## Supplementary data for "3D reconstructions of parasite development and the intracellular niche of the microsporidian pathogen *E. intestinalis*"

### Supplementary figure 1

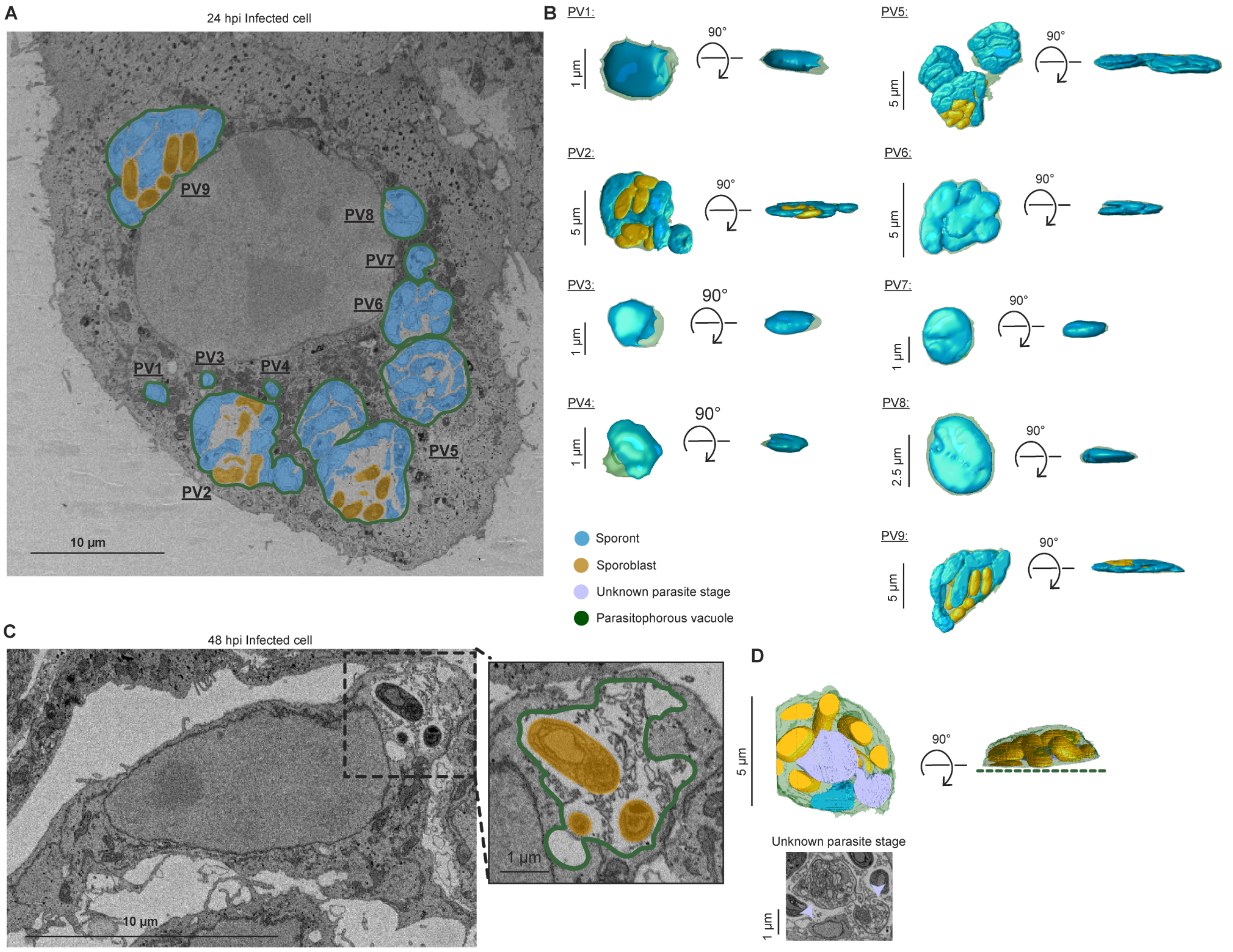


**Supplementary figure 1. SBF-SEM of Vero cells infected with *E. intestinalis*.** (A) SBF-SEM micrograph of an infected cell at 24 hpi. (B) 3D reconstructions of parasitophorous vacuoles from the cell in (A). (C) SBF-SEM micrograph of an infected cell at 48 hpi. (D) 3D reconstruction of the parasitophorous vacuole from the cell in (C) (top), dotted green line indicates the missing region of the parasitophorous vacuole in our dataset, which did not capture the whole cell and a single slice of SBF-SEM data through the parasitophorous vacuole (bottom). Light purple arrowheads indicate an unknown parasite stage.

### Supplementary figure 2

A

| developmental stage | 24-hpi | 48-hpi |
|---|---|---|
| sporont | 48 | 16 |
| sporoblast | 32 | 48 |
| mature spore | - | 140 |
| **total** | 80 | 204 |

B

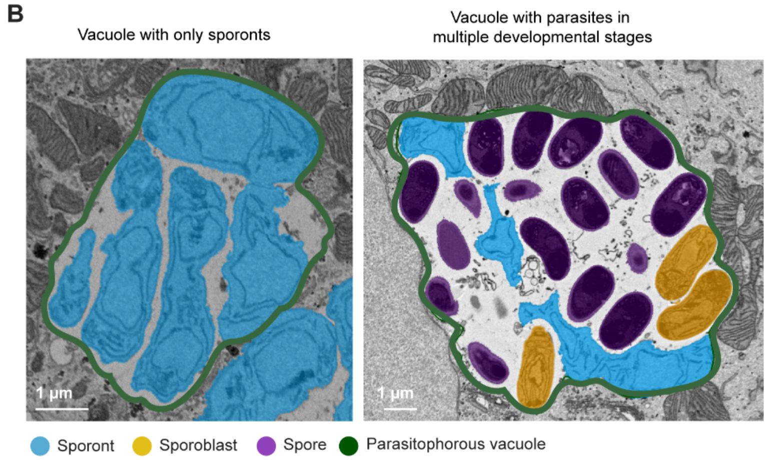


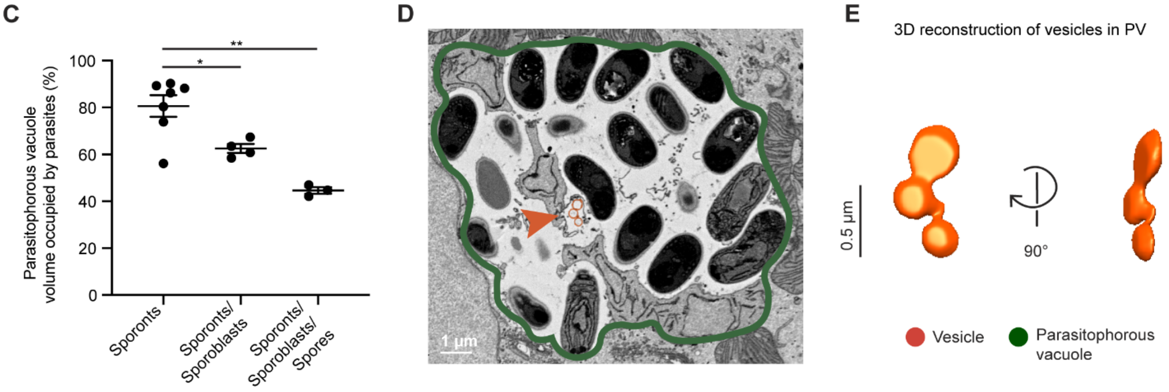


**Supplementary figure 2. Analysis of parasitophorous vacuoles containing *E. intestinalis* from SBF-SEM data of infected cells**. (A) Table of *E. intestinalis* developmental stages observed in 24 hpi and 48 hpi datasets. (B) 2D slices through SBF-SEM data for 24 hpi (left) and 48 hpi (right) for different vacuoles, with segmentation annotation shown. (C) Quantification of the percentage of parasitophorous vacuole occupied by parasites. (D) Single slice of SBF-SEM data through a parasitophorous vacuole, highlighting vesicles (orange) observed in the vacuole. (E) 3D reconstruction of vesicles observed in the parasitophorous vacuole. Mean +/- SEM, *p<0.05, **p < 0.01, calculated from Student's t test for C.

### Supplementary figure 3

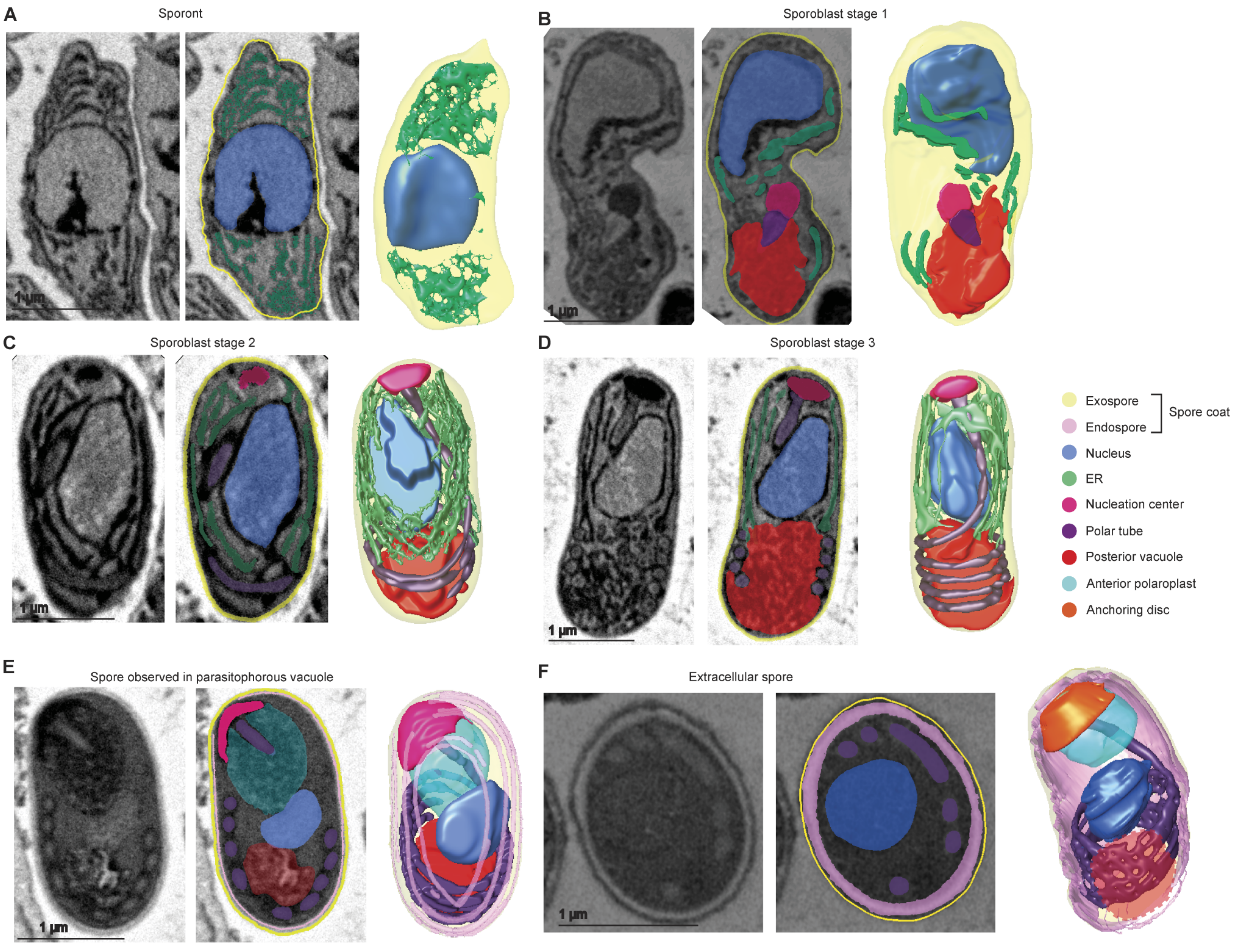


**Supplementary figure 3. 3D reconstructions of *E. intestinalis* parasites at different stages of development.** Representative 2D slices of SBF-SEM data are shown unannotated and annotated, alongside the corresponding 3D reconstruction for a sporont (A), Stage 1 sporoblast (B), Stage 2 sporoblast (C), Stage 3 sporoblast (D), spore (E), all within parasitophorous vacuoles, and an extracellular spore purified from Vero cells (F).

### Supplementary figure 4

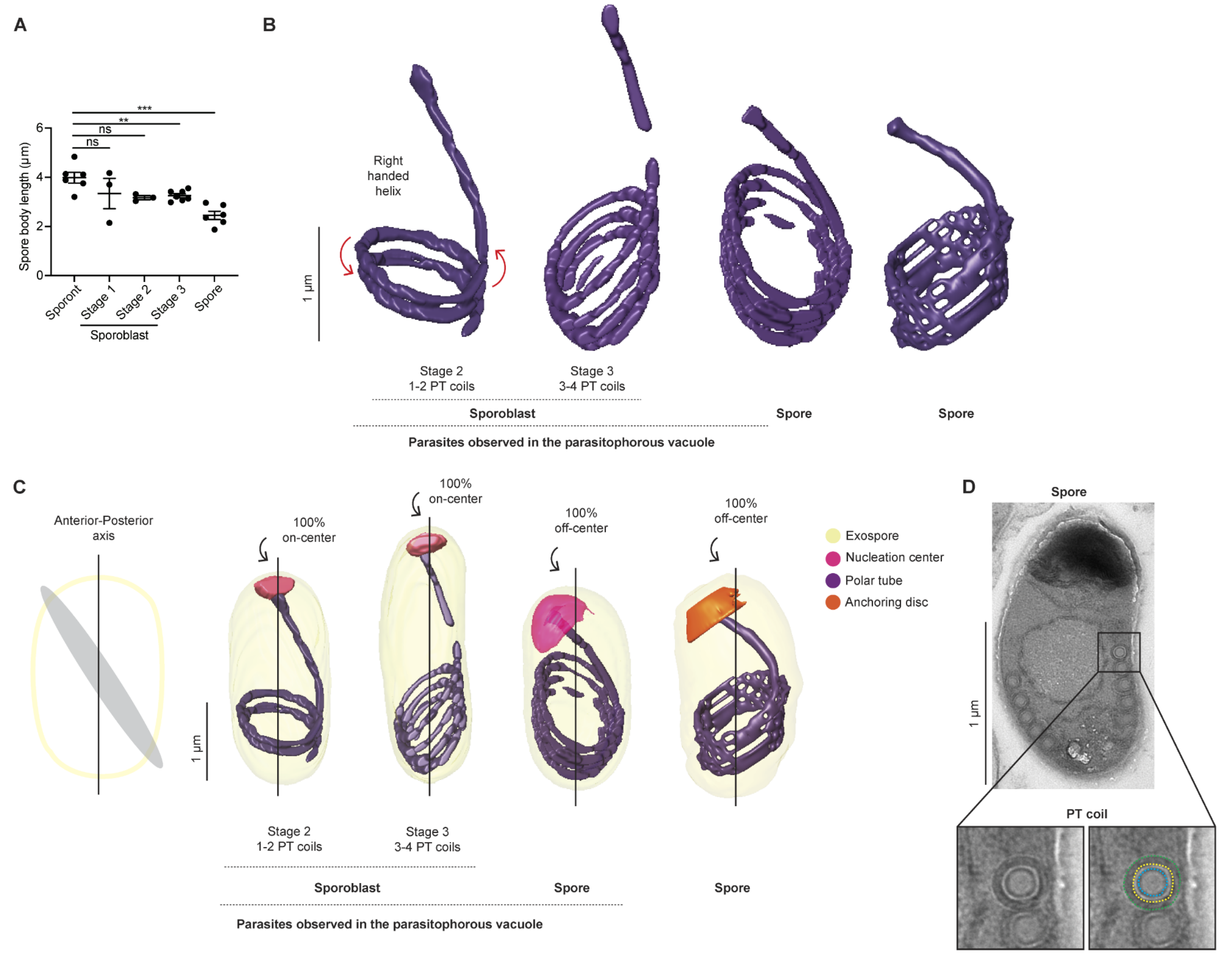


**Supplementary figure 4. Polar tube organization in *E. intestinalis* developmental stages.** (A) Quantification of spore body length at each developmental stage. (B) Representative 3D reconstructions of the polar tube across different stages of development. (C) Position of nucleation center (magenta) or anchoring disc (orange) relative to the anterior-posterior axis. (D) TEM micrograph of spores observed in parasitophorous vacuoles. Each polar tube coil is composed of multiple layers. Mean +/- SEM, $^{**}p < 0.01$, $^{***}p < 0.001$, not significant ($p > 0.05$), calculated from Student's t test for A.

### Supplementary figure 5

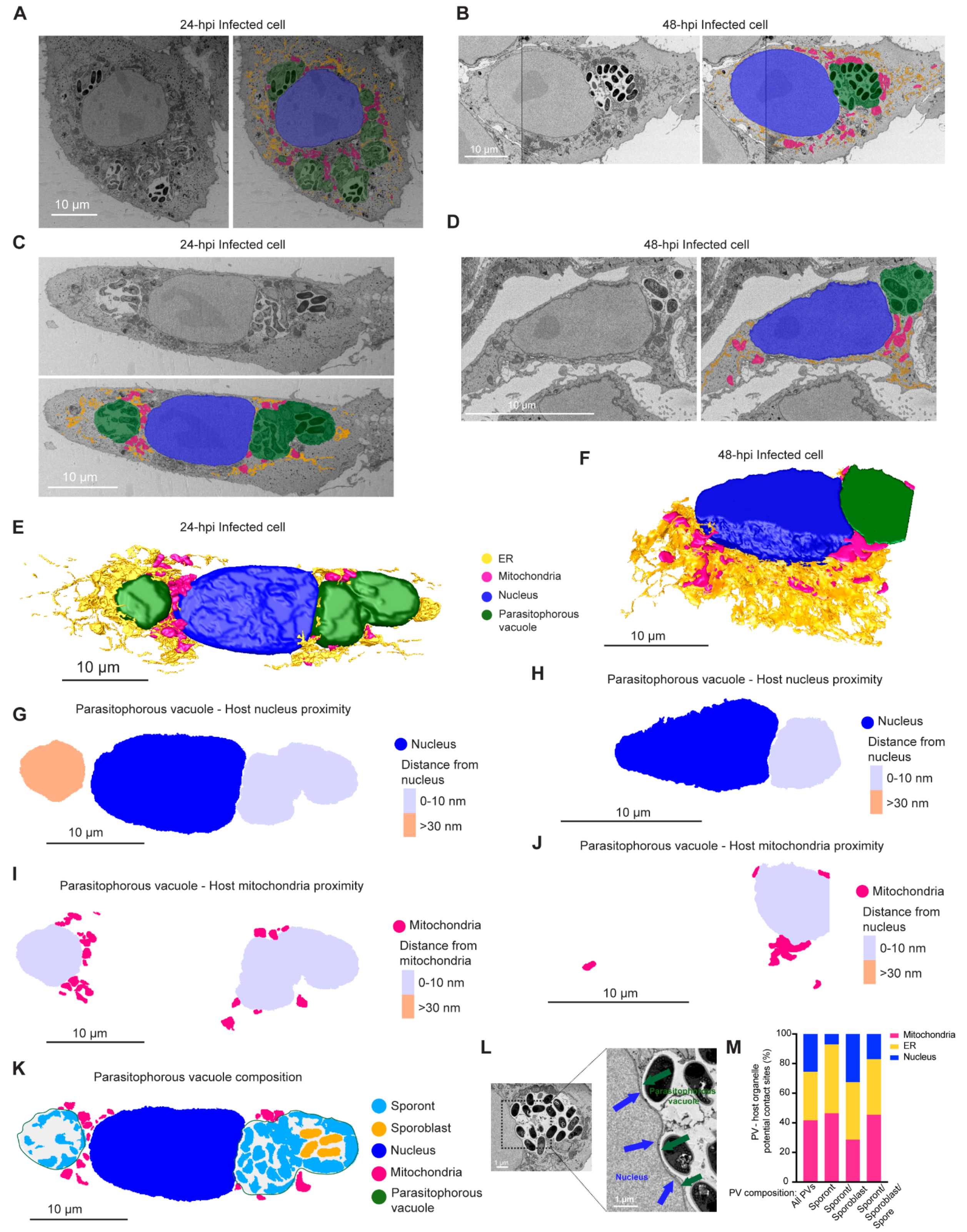


**(Figure legend at top of next page)**

**Supplementary figure 5. 3D reconstructions and analysis of the *E. intestinalis* niche in Vero cells.** (A-D) Single slice from an SBF-SEM dataset of infected cells. Micrographs are shown unannotated and annotated with segmented features side by side for 24 hpi (A, C) or 48 hpi (B, D) cells. (E-F) 3D reconstruction of an infected cell from 24 hpi (E) or 48 hpi (F). (G-H) Distance maps showing the minimal distances from host nucleus to parasitophorous vacuole at 24 hpi (G) and 48 hpi (H). (I-J) Distance maps showing the minimal distances from host mitochondria to parasitophorous vacuoles at 24 hpi (I) and 48 hpi (J). (K) Slice through a 3D reconstruction of the intracellular niche at 24 hpi showing vacuole composition. (L) Slice through SBF-SEM data highlighting indentations in the nuclear membrane (blue arrows) adjacent to the parasitophorous vacuole (green arrows). (M) Graph of the potential contact sites observed between the parasitophorous vacuole and host cell organelles across all vacuoles or separated based on parasite developmental stage.

### Supplementary figure 6

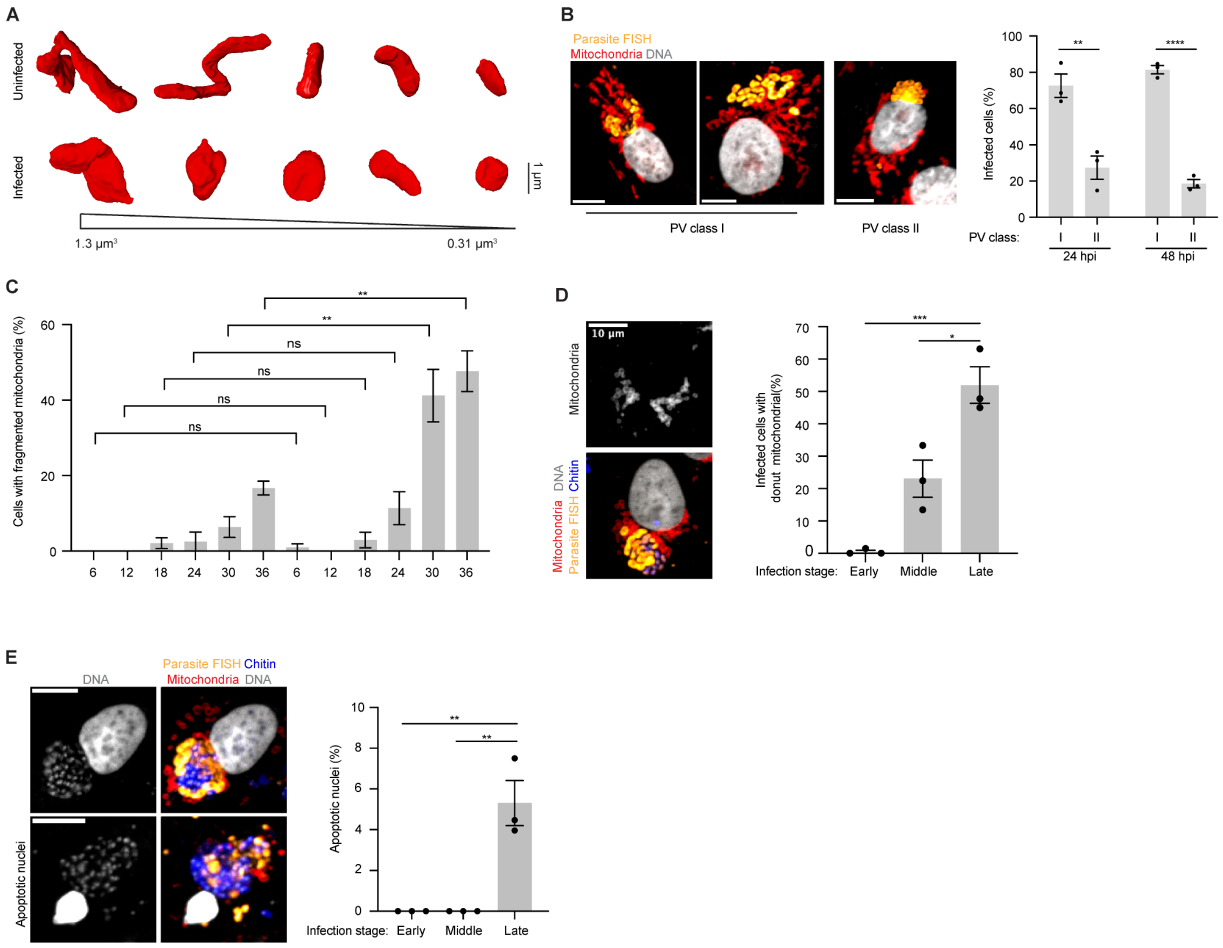


**Supplementary figure 6. Analysis of host mitochondria remodeling in infected cells.** (A) A comparison of mitochondria from uninfected and infected cells, showing the change in mitochondrial morphology observed. (B) Representative micrographs and quantification of parasitophorous vacuole distribution relative to host mitochondrial network in infected Vero cells. Host mitochondria (Tom70 antibody, red), *E. intestinalis* RNA FISH (yellow), DNA (nucblue, gray). Mean +/- SEM from three biological replicates, n=50 cells per experiment. (C) Quantification of mitochondrial fragmentation in uninfected and infected Vero cells for 6 hpi, 12 hpi, 18 hpi, 24 hpi, 30 hpi, and 36 hpi. Cells were fixed and analyzed by RNA FISH and immunofluorescence. Mean +/- SEM from three biological replicates, n=100 cells per experiment. (D) Representative micrograph and quantification of donut mitochondria morphology in early, middle and late stage infected cells. Mean +/- SEM from three biological replicates, n=100 cells per experiment. (E) Representative micrograph and quantification of apoptotic nuclei in early, middle and late stage infected Vero cells. Mean +/- SEM from three biological replicates, n=100 cells per experiment. *p<0.05, **p < 0.01, ***p < 0.001, ****p < 0.0001; ns, not significant (p > 0.05), calculated from Student's t test for B-E.

**Tables**
**Supplementary table 1:** Analysis and composition of parasitophorous vacuoles across all SBF-SEM datasets of infected Vero cells at 24 hpi and 48 hpi.

**Movies**
**Movie 1:** 3D reconstruction of a parasitophorous vacuole from an *E. intestinalis*-infected Vero cell.
Representative reconstruction of a parasitophorous vacuole (green) containing sporonts (blue).

**Movie 2:** 3D reconstruction of an *E. intestinalis* sporont.
Representative reconstruction of an *E. intestinalis* sporont. Each color represents an individual organelle: exospore (yellow), nucleus (blue), ER (green).

**Movie 3:** 3D reconstruction of an *E. intestinalis* Stage 1 sporoblast.
Representative reconstruction of an *E. intestinalis* Stage 1 sporoblast. Each color represents an individual organelle: exospore (yellow), nucleus (blue), ER (green), nucleation center (magenta), polar tube (purple), posterior vacuole (red).

**Movie 4:** 3D reconstruction of an *E. intestinalis* Stage 2 sporoblast.
Representative reconstruction of an *E. intestinalis* Stage 2 sporoblast. Each color represents an individual organelle: exospore (yellow), nucleus (blue), ER (green), nucleation center (magenta), polar tube (purple), posterior vacuole (red).

**Movie 5:** 3D reconstruction of an *E. intestinalis* Stage 3 sporoblast.
Representative reconstruction of an *E. intestinalis* Stage 3 sporoblast. Each color represents an individual organelle: exospore (yellow), nucleus (blue), ER (green), nucleation center (magenta), polar tube (purple), posterior vacuole (red).

**Movie 6:** 3D reconstruction of an *E. intestinalis* spore observed in the parasitophorous vacuole.
Representative reconstruction of an *E. intestinalis* spore. Each color represents an individual organelle: exospore (yellow), nucleus (blue), ER (green), nucleation center (magenta), polar tube (purple), posterior vacuole (red), anterior polaroplast (teal).

**Movie 7:** 3D reconstruction of an *E. intestinalis* spore purified from Vero cells.
Representative reconstruction of an *E. intestinalis* spore. Each color represents an individual organelle: exospore (yellow), nucleus (blue), ER (green), polar tube (purple), anchoring disc (orange), anterior polaroplast (teal). Posterior vacuole could not be segmented accurately in this spore and is not shown in the 3D reconstruction.

**Movie 8:** Live-cell imaging of host mitochondria remodeling in an *E. intestinalis* infected Vero cell.
Time lapse video of host mitochondria fragmentation (red) in an *E. intestinalis* infected Vero cell corresponding to Figure 6E in the main text.

**Movie 9:** Live-cell imaging of host mitochondria remodeling in an *E. intestinalis* infected Vero cell.
Time lapse video of host mitochondria fragmentation (red) in an *E. intestinalis* infected Vero cell that undergoes cell division.

**Movie 10:** *E. intestinalis* life-cycle.
The movie incorporates our findings in the context of what is known from the literature.

**Movie 11:** Development of an *E. intestinalis* parasite.
The movie incorporates cell shape and size changes, as well as the development of individual organelles that could be annotated in SBF-SEM datasets.

**Movie 12:** Model of *E. intestinalis* polar tube development.
Model of polar tube development based on data from SBF-SEM reconstructions.

**Movie 13:** Modeling polar tube coiling.
A stiff tubing (analogous to the polar tube) coils within the confines of a bottle (analogous to the spore) when continuously threaded through the top.
